# Networks of Phenological Synchrony Reveal a Highly Interconnected Ecosystem and Potential Vulnerability to Climate-Driven Mismatches

**DOI:** 10.1101/2022.07.11.499341

**Authors:** Alexis Garretson, Natalie Feldsine, Megan Napoli, Elizabeth C Long, Rebecca E Forkner

## Abstract

As anthropogenic climate change alters species’ phenology, phenological shifts may cascade to disrupt species interactions to impact ecosystem functioning. We present a 108-year phenology dataset of 8,840 event dates for 251 phenophases for seven amphibian species, 58 birds, 14 insects, and 163 plant species, including 52 species introduced to New York. The dataset was collected at a single location in the Northeastern United States, providing continuity in monitoring since the early 1900s. We show that linear phenology analyses can underestimate the magnitude of phenological shift relative to circular methods, particularly for species experiencing extreme advancements. However, species phenologies are generally advancing, with faster advancements of insects and amphibians compared to birds and plants. Additionally, in our dataset, species with event dates later in the year are advancing more rapidly than species earlier in the year, and this relationship is stronger for animals than for plants. We present a novel, network-based approach for visualizing community and ecosystem-scale phenological synchrony. Using this approach, we find a high degree of synchrony between the monitored species, and this approach reveals that plants are more central in the phenological network, as well as species with phenological events earlier in the year. While many synchronous species are shifting at relatively similar rates and display similar temperature sensitivities, we highlight two species interactions potentially vulnerable to changing climate: Eastern Tent Caterpillars and Monarchs. Our results illustrate the utility of long-term ecological monitoring for investigating ecosystem responses to climate change and identifying potentially vulnerable phenological networks.

**Significance Statement:** The purpose of this study is to understand how climate change has affected the phenology of an ecological community for over 100+ years. We present a novel approach to analyzing and visualizing community-level phenological data. We find that plants are central to phenological networks, as are species that flower, fruit, or undergo other phenological events earlier in the year. This is important because understanding which species are most central to an ecosystem, as well as which species are vulnerable to climate-driven mismatches (e.g., a butterfly emerges before the flowers that it feeds on bloom) that could cascade through an ecosystem.

## Introduction

Changes in the timing of species phenological events are among the most sensitive and well-studied ecological responses to anthropogenic climate change. In the Northern Hemisphere, the growing season of plants has lengthened an average of 10–12 days (1–3). However, significant taxonomic and life history-related variability exists in the magnitude of phenological shifts (4–7). Within an ecological community, different species show different magnitudes of responses to climate change and may respond differentially to temperature and precipitation effects (8–11). These differences may lead to disruption of phenological synchrony within ecological communities and can disrupt species interactions (12–15). Recent research has focused on the consequences of this extended growing season for food web processes, such as the synchrony of flowers with pollinators (16), and for ecosystem processes, such as carbon uptake (17).

Some studies suggest that the maintenance of synchrony may be quite common, despite advancing climate change (16, 18, 19). For example, a 20-year forest phenology study by Both et al., 2009 demonstrated a relationship between budburst, caterpillars that consumed the plants, and the hatch dates of passerine birds that consumed the caterpillars. Some evidence suggests that climate change is less likely to disrupt synchrony and lead to a phenological mismatch in lower trophic levels (17, 20, 21). Moreover, antagonistic interactions are expected to be more likely to lead to asynchrony and mismatches than mutualistic interactions (22, 23). Much effort has been invested in determining the synchrony between the mutualistic plant-pollinator appearance, but studies find differing results: while some report either higher advancement of flowering (24) or insects (25, 26), others report similar rates of advancement of both groups (27). Additionally, the majority of studies of synchrony have investigated only a small number of species, and have not provided insights into how networks of interacting species phenologies in an ecosystem may respond to climate variability or how this may translate into broader community change.

Including increasing numbers and diversity of species can introduce statistical challenges, particularly including phenophases that occur near the boundary of the calendar year. This is because phenological analyses are traditionally conducted using only linear analysis models, in regions with a single growing season, and tend to focus on summer or spring phenophases that do not cross the boundary of the yearly date cycle (28–30). However, it is increasingly recognized that the widely observed changes in spring and summer are also seen in fall and winter (15, 31–34). Circular statistics offer a potential solution to concerns with modeling cyclical phenomena with phenological datasets that cover much of the yearly cycle, by expressing dates as angular directions distributed across a circumference (35, 36). While circular statistics are well suited to study phenology, very few studies have used a circular analysis approach (37–39), and using circular approaches may improve the community and ecosystem-level predictions of the impacts of climate change.

Determining the impacts of climate change on community-level phenological change requires long-term monitoring data, including species with diverse taxonomy, ecological roles, and life histories. However, these data are incredibly scarce, and most existing studies of species interactions and floristic communities rely on data collected for alternative purposes (e.g., museum and herbarium specimens) and short-term experimental or observational data. However, researchers at the Mohonk Preserve have been documenting the phenology of plants and animals since 1912 in the Shawangunk Mountains of New York, USA. Compared to existing resources, these data have been systematically collected to monitor phenology with relatively consistent effort and currently stretch more than 100 years. Alongside Mohonk Preserve’s long-term weather monitoring, our study leverages this phenological monitoring data to investigate the impacts of climate change on community-wide plant phenology, species interactions, and community composition. Using these data, we (1) evaluate the differences in shift estimations between linear and circular regression approaches, (2) assess differences in the phenological shift by organismal life history, (3) investigate the relationship between phenological temporal shift and sensitivity to temperature, and (4) synthesize these results to determine impacts of climate change on the Mohonk Preserve ecosystem.

## Results

Phenology data were collected at The Mohonk Preserve, a more than 8,000-acre land trust located in the Shawangunk Mountains in the Hudson Valley of New York. The area has experienced an increase in average precipitation from an average of 1198 mm/year (1931-1989) to an average of 1315 mm/year (1990-2020), and average minimum temperatures have increased approximately 1°C (4.6°C to 5.5°C) over the same period while average maximum temperatures have increased approximately 1.5°C (13.6°C to 15.0°C). The phenological data presented here extend from 1912 to 2020, but the collection remains ongoing. The species includes documentation of 7 amphibian species, 28 long-distance migrating birds, 14 medium-distance migrating birds, 16 short-distance migrating birds, ten butterfly species, four other insects (honey bees, Japanese beetles, katydids, and dog-day cicadas), 69 dicot forbs, 22 monocot forbs, and 72 woody plant species. The resulting dataset consists of 8,840 dates representing 251 phenological events, with an average of 35.2±23.2 (11-92) years of phenology observations, ranging from 1912 to 2020.

Comparing the linear to the circular estimated mean dates showed that, on average, linear mean dates were 0.060±0.21 days later than circular mean dates. However, the dates were highly correlated, with a correlation coefficient of 1 (*P*<0.0001). The maximum difference between the two estimates was 1.15 days (*Vanessa atalanta* adults), and the minimum was - 1.65 (*Colaptes auratus* adults). The differences between linear and circular mean dates were significantly higher for animal phenophase estimates (0.09±0.30) than for plants (0.04±0.15, Wilcoxon *P*=0.0002). We did not find a significant relationship between the difference between the circular and linear estimated mean date and the mean event date itself in these data (*P*=0.11). The estimates of the phenological shift were also highly correlated between the two models (R=0.99, *P*<0.0001). However, the divergence in shifts was higher at values farther from 0 (more positive or negative estimated shifts, Figure 1A). Unlike the estimates of mean phenophase occurrence dates, the mean difference between the absolute values of the shift estimates was -0.02±0.08 days, indicating the circular estimates tended to be of a larger magnitude than the linear estimates. The maximum difference in shifts was 0.03 (*Prunus serotina* fruiting), and the minimum was -0.87 (*Speyeria cybele* adults). Based on these findings, we continued with the circular estimates of shift and mean occurrence dates for the remainder of the paper.

**Figure 1.**
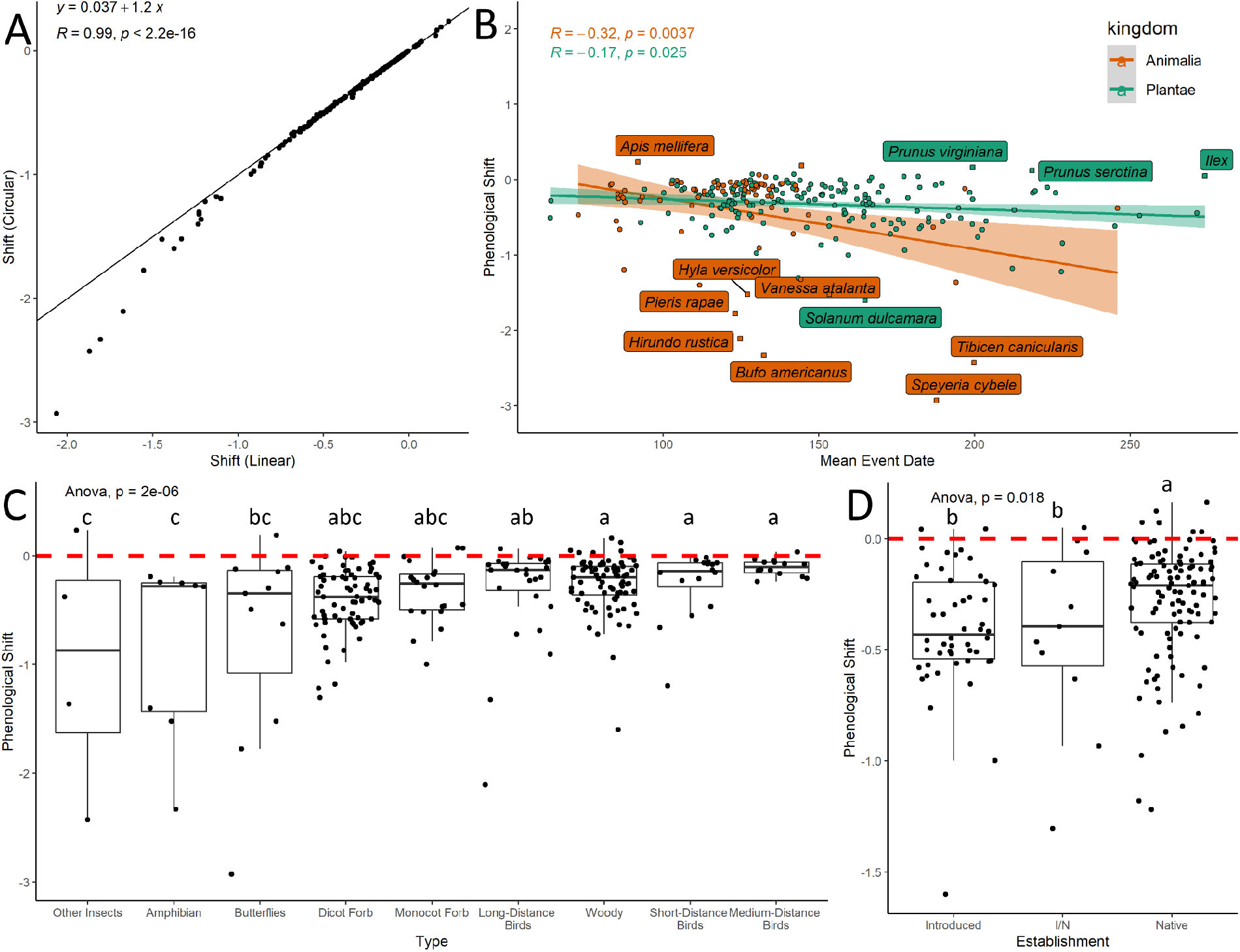
A) Comparison of the linear and circular estimates of the magnitude of the yearly shift in days of the species phenology over time. Each point represents a single estimate of a species phenophase shift, and the line shows the line y=x. B) Correlation between the circular estimate of the mean event date and the circular estimate of the phenological shift. Labeled points show species with extreme phenological shifts, and the blue line shows the regression line for the correlation between these measures. C) Boxplot of the circular phenological shifts of each class of species, each point (jittered to avoid overplotting) represents a single species. The dashed red line indicates a 0 day per year phenological shift, so points above the line are a delay in phenology while the points below the line are an advancement in phenology. D) Boxplot of the circular phenological shifts of the establishment type of each plant species. I/N indicates species or taxa that include both introduced and native members.

The mean shift across all species and phenophases was an advancement of 0.35±0.42 days per year. The largest animal advance observed was for adults of *Speyeria cybele*, which advanced by 1.46 days per year across 25 years of monitoring for a total advancement of 36.5 days since 1993. In comparison, the largest plant delay was in *Prunus virginiana*, which was delayed by 0.08 days per year, while the largest plant advancement was in *Solanum dulcamara* that advanced by 0.80 days per year. The full list of species shifts and mean event dates recovered from our dataset is available as Supplemental Data Sheet 1. We found a negative correlation between the mean phenological event date and the estimated shift for both plants in animals indicating that events occurring later in the year experienced greater advancements than those occurring earlier (*R*=-0.19, *P*=0.002, Figure 1B). However, the correlation was stronger for animals (R=-0.31, *P*=0.003) than for plants (R=-0.17, *P*=0.025). Comparing the shifts by life history groupings was significant (F = 15.04 P<0.0001, Figure 1C), and a Tukey post hoc test showed shifts experienced by other insects and amphibians were significantly lower than woody plants, short-distance migrating birds, and medium-distance migrating birds. While there were no significant differences based on the establishment (native vs. introduced vs. monitored taxa that contain both introduced and native members, I/N) across the whole dataset (*P*=0.16); within the plants, native plants showed significantly lower shifts (−0.14±0.13 days per year) than introduced (−0.20±0.14 days per year) and groups with both introduced and native forms (−0.21±0.21 days year) species (F=4.096, *P*=0.018, Figure 1D).

In general, species shifts were advancements (Figure 2A-G), and temperature sensitivities followed a similar pattern. Temperature sensitivities varied across the monitored species, with an average temperature sensitivity of -2.29*±*2.56, meaning that a one-degree increase in temperature was associated with an average of 2.7-day advancement in the first occurrence date of the species phenophase. The overall magnitude of the temperature sensitivity varied between plants and animals, with plants experiencing higher average delays than animals (animal=-2.11*±*2.42, plant = -3.12*±*2.57. Wilcoxon *P*=0.004). There was a significant relationship between the temperature sensitivity and the phenological shift of a species phenophase, with a higher correlation between these metrics for plants than for animals (Figure 2G). The correlation coefficient between animal temperature sensitivities and the phenological shift was 0.32 (*P*=0.003), while the correlation between plant temperature sensitivities and phenological shifts was 0.6 (*P*<0.0001, Figure X). An ANOVA revealed significant variation in the life history grouping of the species (F=3.43, *P*=0.0009), with amphibians and insects having larger phenological responses to single-degree changes than medium- and long-distance migrating birds (Figure 2H).

**Figure 2.**
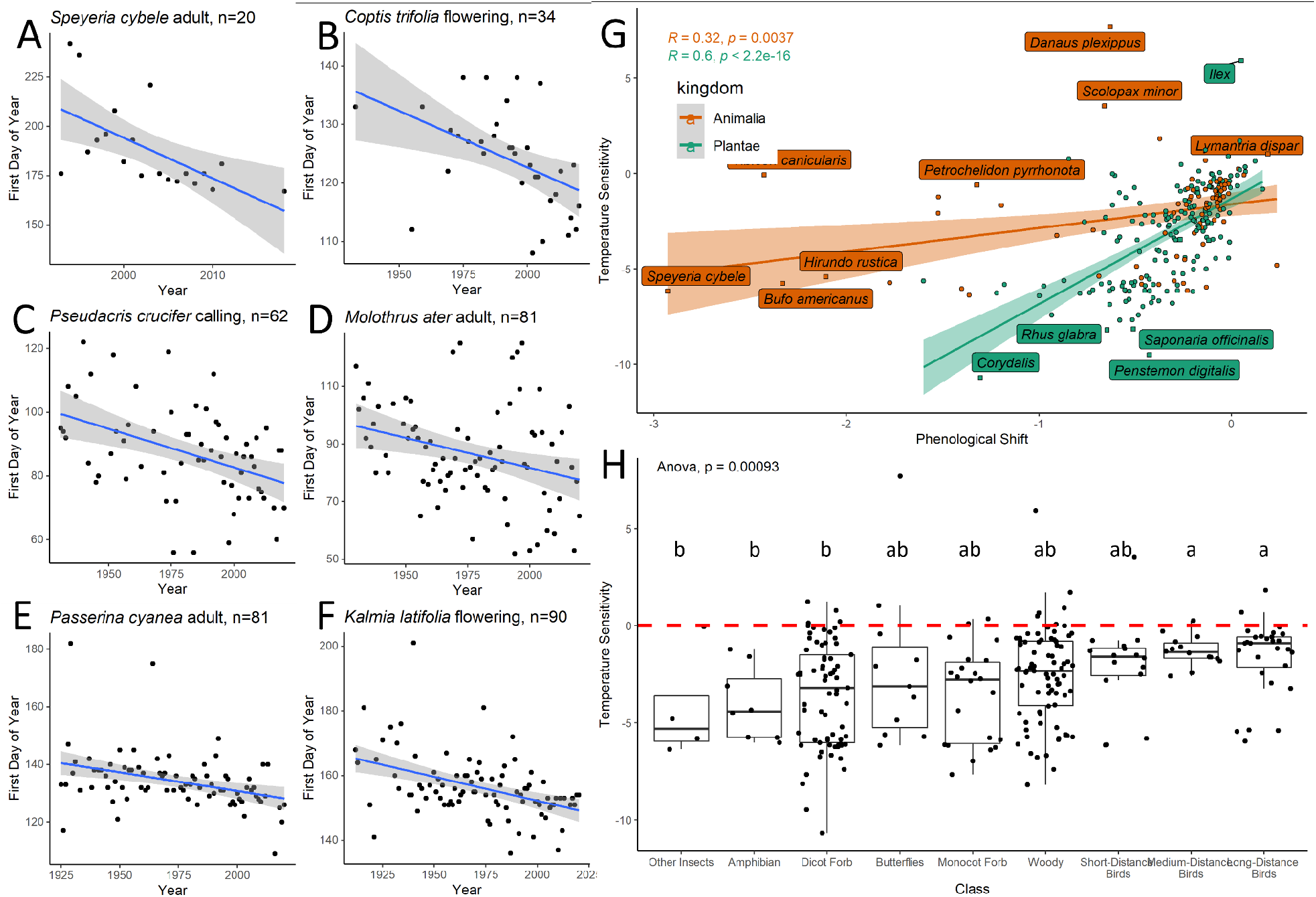
A-F) Scatter plots of the year of observation by the DOY of the first occurrence in that year for various species. G) Scatter plot of the temperature sensitivity of a species against its phenological shift colored by species type, regression lines are best fit linear regressions with a shaded 95% confidence interval. H) Boxplots of the temperature sensitivity of each species grouped by common life-history traits with letters indicating Tukey HSD significant groupings.

In total, 8,467 phenophase pairs that had mean dates within 14 days of each other had sufficient numbers of overlapping years to be analyzed. The mean correlation of these phenophases was 0.24±0.31, indicating generally weak positive synchrony within the dataset. The correlation analyses revealed 352 phenophase pairs occurring within two weeks of each other in high synchrony (R > 0.75), containing 158 unique species. A high degree of correlation between phenological events would suggest that the phenological events may be responding to similar climate or anthropogenic factors or may indicate a causal relationship (e.g., plant-pollinator, insect:host plant, or bird:food source). Many of these species were likely highly synchronous because of similar responses to climate features, for example, synchrony between two flowering plants.

Of the highly synchronous species, 348 phenophase pairs could be arranged into a network of interacting phenophases demonstrating phenological synchrony of the monitored species across the Mohonk Preserve ecosystem (Figure 3A & 3B). In total, four pairs did not link into the extended network and had to be excluded from further analyses. The network included flowering, fruiting, adult, and tents observed phenophases from a total of 152 species, including 21 animals (2 amphibians, 5 butterflies, 9 long-distance birds, 3 medium-distance birds, 2 short-distance birds) and 131 plants (53 dicots forbs, 20 monocot forbs, and 58 woody plants). A subset of identified highly synchronous species likely have causal relationships. For example, the high synchrony between the first *Danaus plexippus* (Monarchs) adult and *Daucus carota* (Queen Anne’s lace) flowering dates likely represents a causal link as adult monarchs feed on Queen Anne’s lace (Figure 3C). Additionally, the host plants of *Malacosoma americanum* (eastern tent caterpillars) are trees in the *Malus* and *Prunus* genera. Therefore, the high synchrony between the dates of tent appearances and the flowering dates of *Malus* spp., *Prunus americana*, and *Prunus avium* likely represent a causal linkage (Figure 3D).

**Figure 3.**
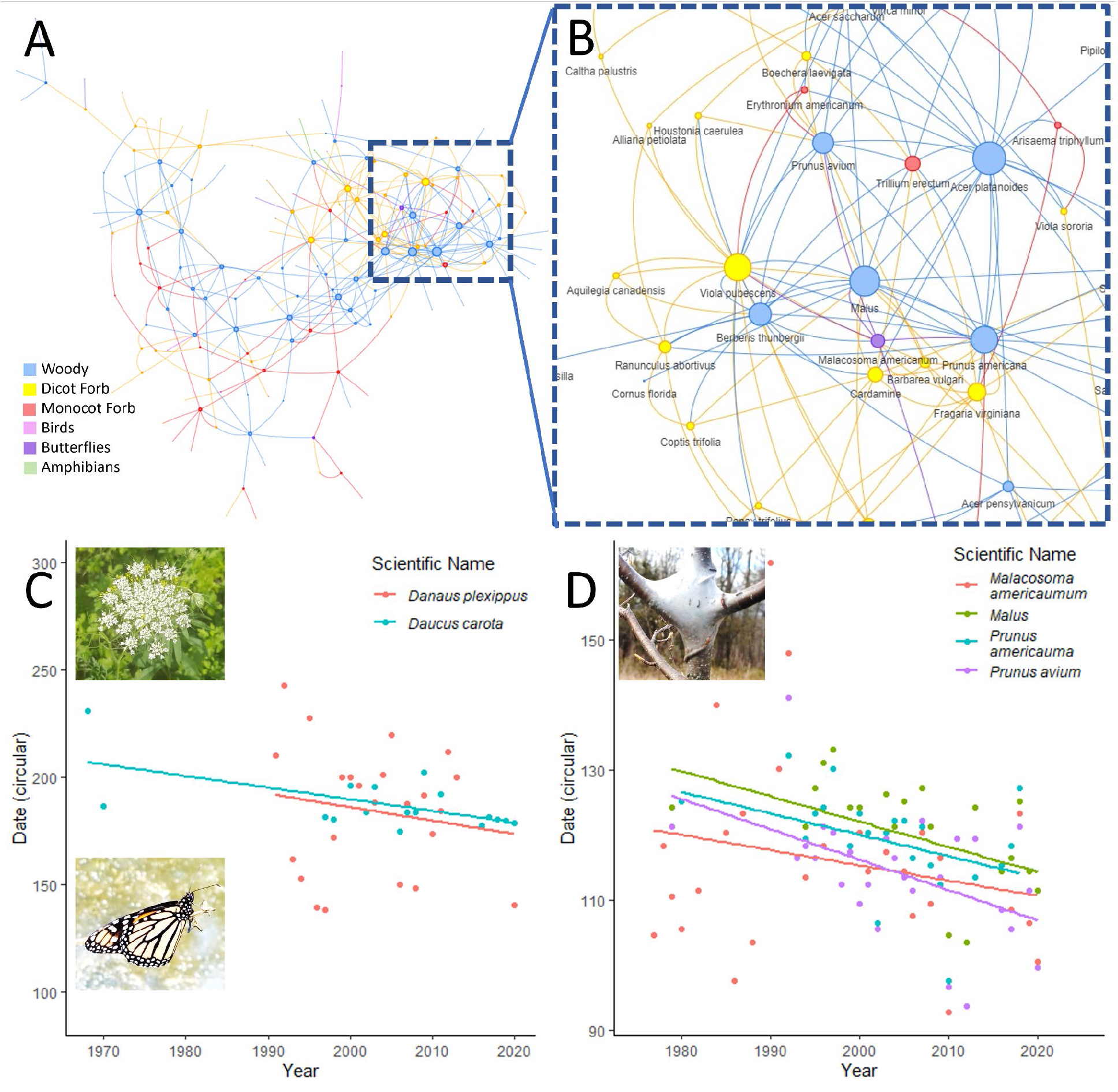
A. Network of the species with highly synchronous phenology in the Mohonk Preserve dataset. Each node is a single species life stage, sized by the number of synchronous interactions. See supplemental file for the full network in HTML format. B. Zoomed in portion of the network (rotated and jittered to highlight species of interest) C. The first event dates for the adult *Danaus plexippus* (monarch) are highly correlated with the first flowering dates for *Daucus carota* (Queen Anne’s lace), which is one of the adult monarch food sources. D. The first events for the *Malacosoma americanum* (eastern tent caterpillar moth) tents, are highly correlated with first flowering dates for *Malus* spp. (apple), *Prunus americana* (American plum), and *P. avium* (sweet cherry).

The most central species in the network were the flowering dates of *Sambucus racemosa* (0.00250), *Rosa multiflora* (0.00249), *Gaylussacia* spp. (0.00246), *Acer platanoides* (0.00245), *Prunus americana* (0.00245), and *Malus* spp (0.00242) (Figure 3B). There was a moderate positive correlation between the centrality of a species and the phenological shift (R=0.3, *P*=0.0001, Figure 4A), with species with lower phenological shifts having higher network centrality. This correlation was stronger for animals (*R*=0.54, *P*=0.012) and weaker for plants (R=0.21, *P*=0.016, Figure 4A). There was also a significant negative relationship between centrality and mean phenological event date (R=-0.38, *P*<0.0001, Figure 4B), with species with earlier event dates having higher centrality in the phenology network (Figure 4B). The correlation between animal centrality and mean event dates was -0.74 (*P*=0.0001), while the correlation between plant centrality and mean event dates was -0.41 (*P*<0.0001). The centrality of the plant phenophases was significantly higher than the centrality of the animal phenophases (Wilcoxon *P*=0.0014, Figure 4C), but there were no significant differences between introduced, introduced/native, and native species (Kruskal-Wallis *P*=0.66). An ANOVA revealed significant differences between life-history categories (F=2.78, *P*=0.009), with woody plants significantly more central than butterflies and long-distance migrating birds (Figure 4D).

**Figure 4.**
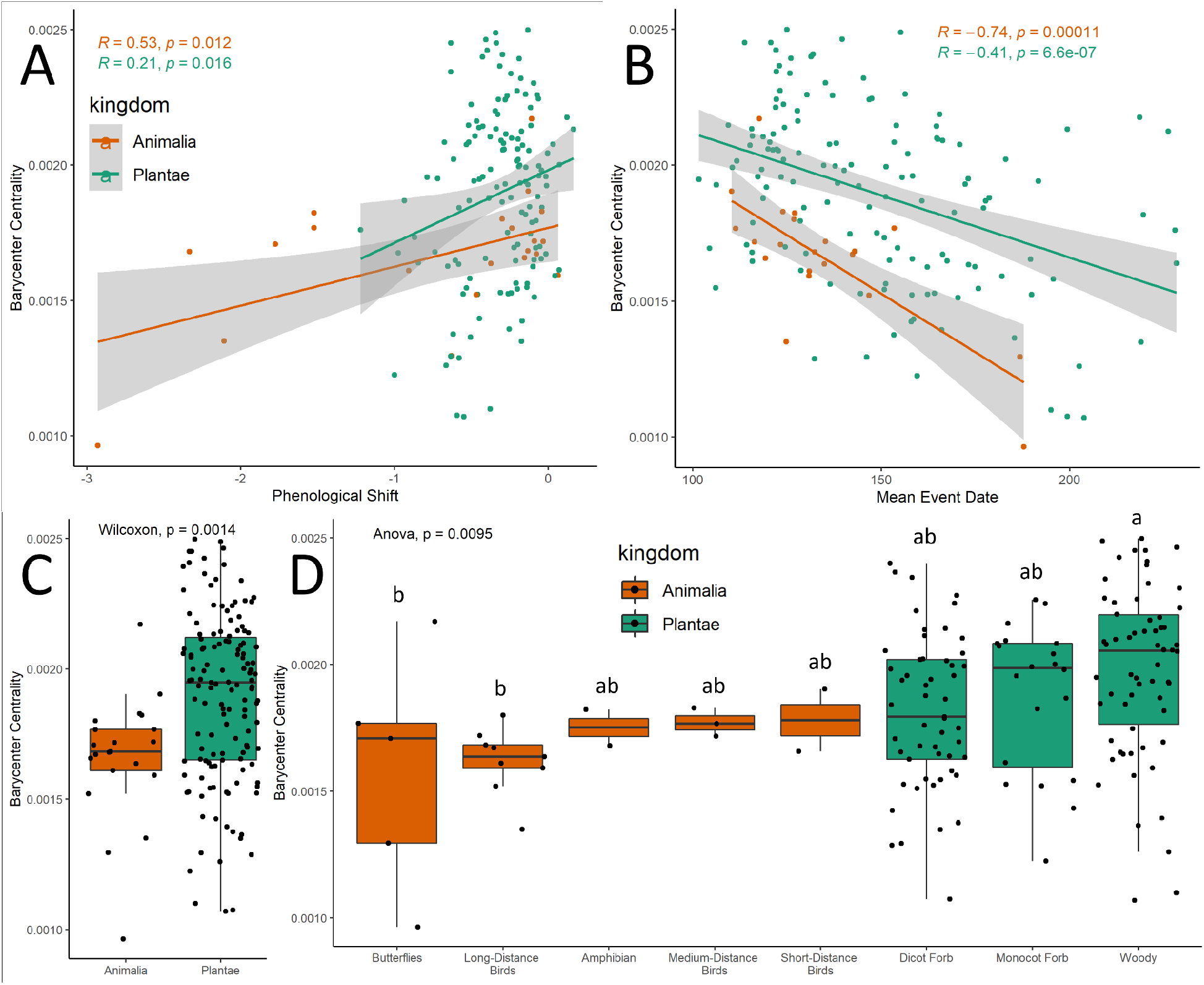
A. Scatterplot of the Barycenter Centrality of a phenophase against the phenophase mean event date and B. the phenological shift. Each point represents a single phenophase in the phenological synchrony network. The line is the best fit linear regression line with the grey area representing a 95% confidence interval. The Spearman’s correlation coefficients and the *P*-values are shown in the upper right. C. Boxplots of the centralities of different species groups in the network and their calculated Barycenter Centrality by kingdom, D. Boxplots of the centralities of different species groups in the network and their calculated Barycenter Centrality by common taxonomic or life-history traits.

Across all species pairs, the mean difference in temperature sensitivities was 0.05±3.22. A total of 17 between-taxa species pairs were both highly correlated (>0.75) but had differences in temperature sensitivities greater than one standard deviation (>3.22) (Supplemental Table 2). On average, the plant species had a higher temperature sensitivity than the correlated animal species (−2.8±5.06). The largest sensitivity difference was between Queen Anne’s lace flowering dates and monarch first adult dates, where each one-degree increase in temperature was associated with a 13-day advancement of the Queen Anne’s lace flowering dates relative to first Monarch adult dates. Similarly, a one-degree increase in the temperature was associated with a 4.09 and 3.65-day advancement of *Malus* sp. and *Prunus avium* flowering, respectively, relative to the first tent date of *Malacosoma americanum*.

## Discussion

In this work, we present a novel, network-based method for analyzing multi-species, long-term phenological data and interrogate this network to uncover potential vulnerabilities to climate-driven mismatch. Our analyses show that species in the Mohonk Preserve ecological community have shifted at varying rates, and these variations follow taxonomic and temporal patterns (Figure 1, Figure 2A-F). Specifically, while we found a general advancement in phenology, we find that insects and amphibians are shifting faster than birds and plants (Figure 1C). This finding is congruent with previous cross-taxa studies, but with our extensive dataset in a single locality, these results add important confirmation to these findings (26, 42, 43). Additionally, we found that native plants have shifted less than introduced plants (Figure 1D). Previous work has found that invasive plants often have higher phenotypic and phenological plasticity (44, 45), which can enable range expansion and competitive advantage over native plants (46). This variability in climate adaptiveness can limit native plant success, which may have cascading effects on ecosystems and interacting species (47).

We also found that species phenology occurring later in the monitoring season are advancing more rapidly, particularly within the monitored animals (Figure 1B). While our phenological monitoring does not span the full year and focuses on spring and summer phenophases, our finding underscores the importance of monitoring species throughout the year and suggests that late summer and fall phenophases may be particularly important in understanding the impacts of climate change on ecosystems and species interactions (31–34). Finally, our results underscore the importance of using appropriate modeling approaches, as linear estimation methods yield lower shifts relative to circular estimation methods, particularly for species that have seen dramatic shifts over the study period (Figure 1A).

By investigating the phenological responsiveness to interannual temperature variability, we found a positive correlation between temperature sensitivity and observed phenological shifts (Figure 2G), however, we find this correlation is stronger for plants than for animals, suggesting that while the majority of plant phenology shifts are temperature-related, animal phenology may be reacting to other anthropogenic impacts or shifts in interacting species. We also find that the taxonomic patterns of differences in phenological shifts were largely maintained in the temperature sensitivities, with insects and amphibians showing greater sensitivity (Figure 2H). Together, these results highlight the importance of paired climate and phenology monitoring in single localities, and the importance of developing a strong understanding of the diverse drivers of ecosystem change.

Evaluating the synchrony of phenological events in this dataset revealed a highly interconnected phenological network (Figure 3A, B). High phenological synchrony may not indicate a direct link between two species but could indicate a mediated link through a species not monitored in this dataset. For example, high synchrony between an insectivorous bird and a flowering plant may be synchrony mediated through an unmonitored third species, the insect food species of the bird that uses the flowering plant as a host. Alternatively, it may indicate shared responsiveness to climate variability. While our current analyses do not detangle these various potential causes of synchrony, we are able to identify multiple synchronous species with putative causal, including between adult Monarchs and a host plant (Queen Anne’s Lace) and between Eastern Tent Caterpillar tent dates and host tree phenology (Figure 3C, D).

Additionally, we generally found that synchronous species had similar responsiveness to temperature. However, we did find that the Monarch:Queen Anne’s Lace synchrony had significantly different temperature responsiveness, as did the Eastern Tent Caterpillar Moths:Host trees. This suggests that accelerating climate change may disrupt these two species’ phenological interconnectedness. Eastern tent caterpillars support a variety of parasitoids (48) and are food items for a variety of bird species (49). Phenological shifts in this species, and potential mismatches, may be accelerated by warming (50). Similarly, the Monarch butterfly is a well-studied, charismatic species, and its recent population declines have sparked a number of new studies, and some of the declines have been attributed to reduced breeding-season habitat availability (51). However, both insect species have more than one host plant, which may mean that despite the potential for mismatch for these particular synchronous interactions, the insects may be buffered from severe impacts by being able to switch between available hosts.

Within the network of synchronous phenology at Mohonk Preserve, there were patterns in species interconnectedness. First, we find a relationship between centrality and phenological shift with lower shifts more central to the network (Figure 4A). This follows the observation that plant phenology was more central in the network (Figure 4C), but the relationship between shift and centrality held for plants as well as animals. Similar to previous findings of ecological synchrony (29), this suggests that networks of synchronous phenological may be buffered by a modest shifting of the most central species. We also found that the phenological event date was related to the network centrality, with species with earlier event dates having higher centrality in the phenology network (Figure 4B). Finally, the centrality of the plant phenophases was significantly higher than the centrality of the animal phenophases, and woody plants had the highest centrality (Figure 4C, D). Together, our work demonstrates the utility of single-site, long-term monitoring of phenology for unraveling potential ecosystem-wide climate impacts on phenology.

## Methods

Following the digitization of the Mohonk Preserve archives, these data were made publicly available for research usage (edi.1152.1 https://doi.org/10.6073/pasta/cbdceb7c5be7430a9e653e3f9379a9e8 and GBIF occurrence dataset https://doi.org/10.15468/b6zrtg) (40, 41). While the full phenology dataset is extensive, we only analyzed species-phenophase pairs with more than ten years of data for insects, birds, amphibians, and plants. Phenophases documented included eggs and egg masses, larvae, juveniles, adults, calling adults, flowering, fruiting, and tents observed for Malacosoma americanum. Additionally, the dataset includes 52 species introduced to New York, including three insects.

We converted linear dates (0-365) to degree dates by mapping the linear day of the year (DOY) into 360 degrees, (d * 360) / 365. Leap years were not considered separately, but as no phenological events in any year occurred on December 31st, no DOY 366 was included, and all dates in leap years were shifted forward by one following the leap day to accommodate its presence in the yearly cycle. Then, we found the positive mean circular angle of the event dates by treating each observation as a unit vector or point on the unit circle, converting the angle of the direction of the resultant vector back to a DOY. This approach, in theory, will provide more intuitive results for phenophases spanning the yearly transition, as the mean occurrence date for a phenophase occurring on December 31st (DOY 365) and January 2nd (DOY 2) should be January 1st (DOY 1) not July 2nd (DOY 183.5). We then compared the estimates of the mean event dates derived from the circular and linear DOY approximations and determined whether there were phenophase or taxonomic predictors of significant differences between the two values.

We also investigated the impacts of modeling phenological events with linear vs. circular models. In the linear models, we fit the linear regression DOY = α + β_linear_(t − 1980) + εt, which models the occurrence DOY as a function of the year (t), and includes an estimate of the mean date of the phenological event at the year 1980 (α) and the slope of the change of the date over the time (β), hereafter referred to as the phenological shift or shift. In the circular models, we fit a circular-linear regression in the form of the homoscedastic version of the maximum likelihood regression model proposed by Fisher and Lee (1992). The regression model used the circular date as the angular response and the centered year as the linear predictor. The estimated μ --or the circular mean--is used as an estimate of the phenological mean event date akin to α and the circular coefficient alpha is the coefficient of the fitted model, multiplied by two (β_circular_). We compared the shift estimates derived from the linear models (β_linear_) to the phenological shift estimates from the circular models (β_circular_) to determine whether linear or circular models provided systematic differences in the estimations of phenological shifts for particular taxa, groupings, or phenological events. Using the estimated phenological shifts, we assessed systematic differences in the magnitudes and directions of the shifts related to taxonomic, life history, and establishment grouping using ANOVAs with Tukey posthoc tests and correlation coefficients.

We then fit circular-linear models with the DOY of the phenophase onset as the circular outcome and the average temperature as a linear predictor. This provided a measure of the temperature sensitivity of the phenophase. We used linear regressions to relate the climate sensitivities to the overall phenological shift estimated from the circular regressions and the mean circular event date. We then used ANOVAs to explore whether the variation in climate sensitivities was related to the species taxonomy.

Finally, we explored the synchrony between species phenology as the correlation between the circular occurrence dates of a species’ life-history event. We restricted the set of potentially synchronous events as pairs of events occurring within a two-week window from each other and excluded pairs of life-history events from the same species.Next, we created a representation of community phenological synchrony creating a network of interacting species whose phenology was correlated at a threshold of at least 0.75. We assessed the centrality of each species in the resulting community phenology networks using the Barycenter centrality score calculated as 1 / (total distance from the target node to all other nodes) and determined whether any species with the highest levels of centrality had shift metrics or divergence metrics that could indicate risk of phenological cascade through the network.

## Data Availability

Data are available on GBIF as an occurrence dataset https://doi.org/10.15468/b6zrtg and EDI under package identifier edi.1152.1 at https://doi.org/10.6073/pasta/cbdceb7c5be7430a9e653e3f9379a9e8

## Acknowledgments

We thank all historical data collectors and digitization volunteers for their work and contributions to citizen science, particularly D. Smiley and P. Huth.

## Author Contributions

NF, EL, and MN collected the bulk of the recent data. AG led the data cleaning and preparation, assisted by NF, EL, and MN. AG led the analysis design and implementation, assisted by RF. AG led the writing of the manuscript and figure development. All authors edited and approved the final manuscript.

